# Structome-TM: Complementing dataset assembly for structural phylogenetics by addressing size-based biases

**DOI:** 10.1101/2025.02.08.637224

**Authors:** Ashar J. Malik, David B. Ascher

**Author notes:** Correspondence to Ashar J. Malik, David B. Ascher.

## Abstract

Harnessing the explosion of protein structure data to uncover deep evolutionary relationships requires effective comparison methods. While widely used global alignment techniques are powerful, they can fail to identify homologous structures that differ significantly in size or domain architecture. To address this limitation, we introduce Structome-TM, a web resource for assembling datasets for distance-based phylogenetic reconstruction. Making use of the TM-score to prioritize local structural similarity, Structome-TM excels at identifying these otherwise obscured relationships, allowing users to build a comprehensive structural neighbourhood of proteins suitable for comparison. To facilitate this dataset assembly, the resource accepts both RCSB PDB identifiers and protein sequences as inputs. When querying using a protein sequence, protein structures are predicted in real-time and their respective neighbourhoods determined, enabling analysis where experimentally determined structures may not be available. Through its user-friendly interface, Structome-TM provides a powerful and necessary approach for a more comprehensive exploration of protein evolution. This resource is freely available at: https://biosig.lab.uq.edu.au/structome_tm/.

## Introduction

The large-scale availability of protein structure data has revolutionized the exploration of deep phylogenetic relationships [1, 2]. These relationships are typically determined in a two-part process involving: (1) gathering structures to define the taxon set and (2) comparing these structures to infer evolutionary relationships. The Structome suite is intended to streamline this process. The first tool developed in this suite, Structome-Q (originally published as Structome [3]), uses the Q-score metric from GESAMT [4] to assess global structural similarity. It efficiently identifies related proteins and performs the necessary pairwise comparisons to construct a neighbour-joining tree.

The Q-score metric is defined as:

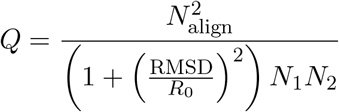

where *N*_align_ is the number of aligned residue pairs, RMSD is the root-mean-square deviation of the alignment, *R*_0_ is a scaling parameter (set to 3.0 Å), and *N*_1_ and *N*_2_ are the amino acid counts of the two structures.

The normalisation by the product of the protein lengths (*N*_1_*N*_2_) makes the Q-score a robust measure of global structural similarity. However, this inherently penalizes alignments where structures share only localized regions of similarity, such as proteins with partial domain matches. As a result, these biologically significant homologues may rank poorly and could be overlooked.

In contrast, the TM-score [5] adopts a different approach:

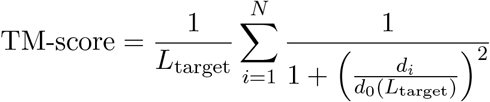

where *L*_target_ is the length of the target protein, *N* is the number of aligned amino acid, *d*_*i*_ is the distance between aligned amino acid pairs, and

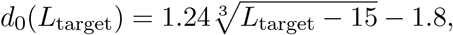

The implementation of this metric within Foldseek [6] prioritizes local similarity by normalizing for alignment length rather than overall protein size. This is particularly advantageous for identifying meaningful alignments between multi-domain proteins that may only share a single domain with the query, highlighting relationships underestimated by the Q-score. The latest resource in this suite, Structome-TM, therefore provides this powerful local alignment methodology. By offering complementary search strategies, the Structome suite provides researchers with a more comprehensive and nuanced toolkit for structural phylogenetics. To clarify the distinctions between these web resources, their features are summarized in Table 1.

**Table 1:** Comparison of Structome-Q with Structome-TM. The table highlights the key differences in methodology and functionality between the original published tool, the updated Structome-Q, and the new Structome-TM.

| Feature | Structome (Original) | Structome-Q | Structome-TM (This work) |
| --- | --- | --- | --- |
| Primary Metric | Q-score | Q-score | **TM-score** |
| Core Engine | GESAMT | GESAMT | **Foldseek** |
| Alignment Focus | Global Similarity | Global Similarity | **Local Similarity** |
| Input Type | PDB ID | PDB ID | **PDB ID & Protein Sequence** |
| Taxon set | Pre-computed (top 50) | User defined | User defined |
| Status | Superseded | Active | **New** |

## Methods

The architecture of Structome-TM builds upon the framework established by Structome-Q [3]. The methodology can be divided into three main components: dataset construction, the search algorithm, and the web server implementation.

### Dataset Construction

The foundation of Structome-TM is a representative, non-redundant set of protein structures derived from the RCSB Protein Data Bank [7] (PDB; www.rcsb.org). To curate this dataset, all PDB entries were first filtered to remove short peptides, retaining only proteins longer than 50 amino acids. This collection was then clustered at 90% sequence identity using USEARCH [8]. Each resulting cluster is represented by a single centroid structure, which serves as a proxy for all its members. This approach makes the subsequent all-versus-all comparison computationally tractable.

### Structural Search and Analysis

To enable rapid querying, an all-versus-all pairwise comparison of all centroids in the database was performed using Foldseek [6], with the resulting TM-scores stored. When a user submits a PDB structure as a query, it is first mapped to its pre-calculated cluster centroid for which the pre-computed results are returned.

To broaden accessibility, Structome-TM also accepts a protein sequence. A submitted sequence is folded into a 3D model in real-time using ESMFold [9]. The resulting predicted structure is then used as the query against the full RCSB PDB. In addition to returning hits, a dedicated column is populated informing if the target protein is a member of the core Structome-TM dataset. Additionally, results for both structure and sequence-based query are annotated with data from several key resources, including SCOP [10], CATH [11], ECOD [12], and the NCBI taxonomy database [13].

### Web Server Implementation

The web application is built using the Flask Python framework and is deployed within Docker containers managed by an Nginx web server. The front-end is developed with standard HTML, CSS, and JavaScript, utilizing AJAX. Interactive 3D visualization of protein structures are rendered using the Mol* viewer [14]. For phylogenetic analysis, neighbour-joining trees are generated from the “1 - TM-score” distance metric using Biopython and are rendered as interactive diagrams in the browser via the D3.js library.

### Usage

Structome-TM offers two query modes, structure-based and sequence-based, each tailored for different input-type, but both leading to a common interface for interactive analysis and phylogenetic tree generation.

### Structure-based Query

When a PDB identifier is submitted, the search is performed against the curated set of Structome-TM centroids. Results are displayed in a sortable table listing hits with a TM-score *>* 0.1. Clicking any row expands it to show the full list of PDB IDs within that cluster and displays an interactive 3D superposition of the query structure against the hit centroid using the Mol* visualiser.

### Sequence-based Query

For protein sequence submissions, the resource first predicts the query’s structure and then searches it against the complete PDB dataset (as of June 2025). The resulting table includes an additional, sortable column with a binary marker. This marker indicates whether a given hit is also a representative centroid within the core Structome-TM dataset.

### Statistics and Analysis

For both query modes, an accompanying histogram summarizes the distribution of TM-scores, providing a rapid overview of result quality. Users can then select hits via checkboxes to assemble a custom dataset for phylogenetic analysis. A key difference between the two modes is the composition of this dataset: the structure-based search allows only curated centroids to be used for tree reconstruction, whereas the sequence-based search allows any target protein to be included, regardless of its centroid status.

Upon submission, a neighbour-joining tree is generated using “1 - TM-score” as the distance metric. The final tree is rendered as an interactive diagram, allowing inspection of individual leaf labels and download in Newick format for further annotation.

### Case Study: Identifying Homology in Multi-Domain Proteins

To demonstrate how Structome-TM captures meaningful alignments missed by global methods, hemoglobin from *Anser indicus* (a single globin domain) was compared with flavohemoglobin from *Saccharomyces cerevisiae* (Figure 1). Flavohemoglobin is a larger, multi-domain protein, comprising a globin domain fused to an FAD-binding domain [15].

**Figure 1.**
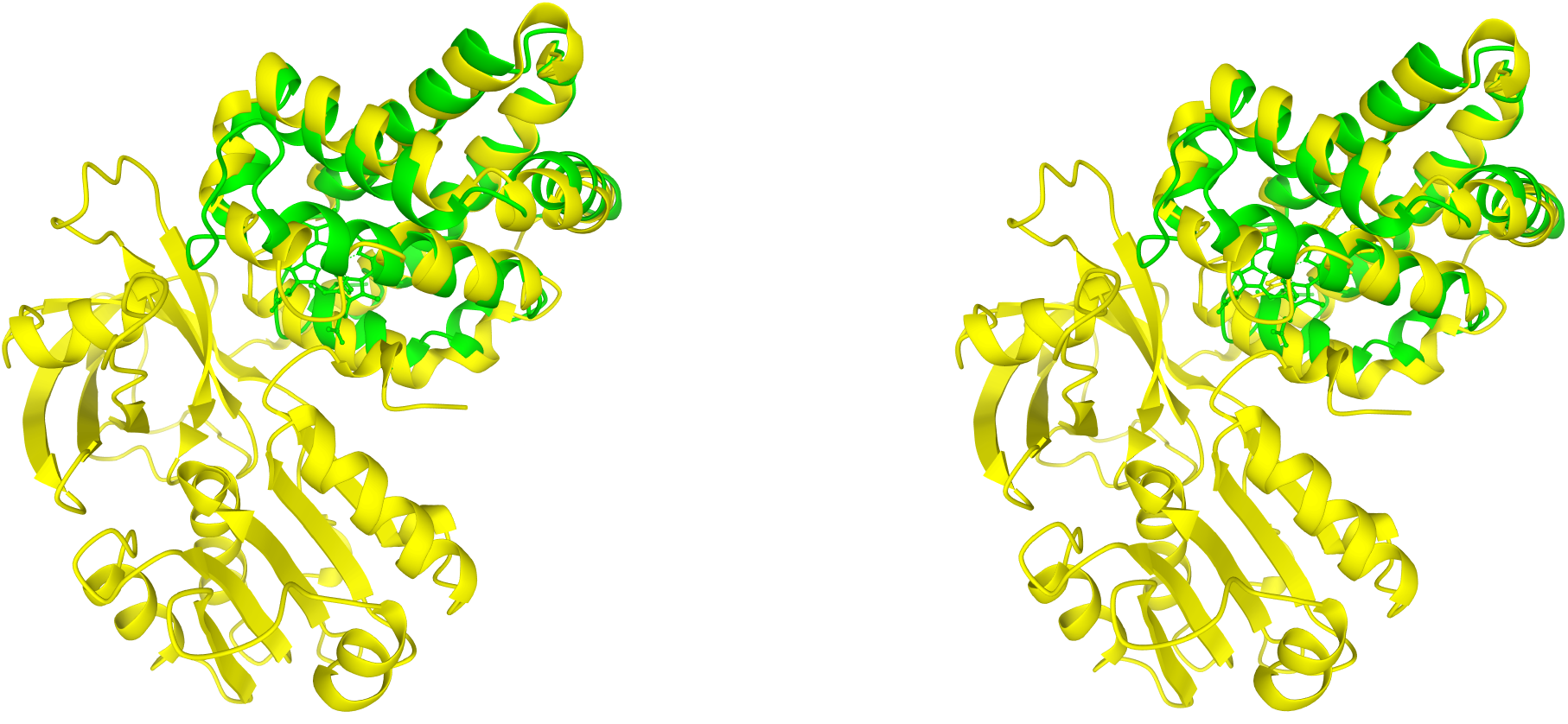
Protein structure alignment between hemoglobin from *Anser indicus* (RCSB PDB accession 1a4f, chain A, 141 amino acids, shown in green) and flavohemoglobin from *Saccharomyces cerevisiae* (RCSB PDB accession 4g1b, chain A, 399 amino acids, shown in yellow). The left panel illustrates the alignment from Structome-Q using the Q-score metric, while the right panel shows the alignment from Structome-TM using TM-score. The Q-score for this alignment is 0.213, whereas the TM-score is 0.803. The lower similarity score from Q-score reflects its global normalization accounting for overall protein sizes. A score of 0.213 indicates that this hit will probably be lost in the background noise.

Due to this size mismatch, the global alignment approach assigns a low Q-score of 0.213. This score is low enough to be indistinguishable from background noise, meaning this significant homologous relationship would likely be missed.

In contrast, by focusing on the shared local similarity, a high TM-score of 0.803 is obtained. This result demonstrates how Structome-TM can successfully build a more comprehensive structural neighbourhood. It ensures that valuable homologous relationships within multi-domain proteins are correctly identified, enabling more complete and accurate downstream phylogenetic analyses.

### Outlook

Structome-Q and Structome-TM are designed to streamline the initial, and often most challenging, step of structural phylogenetics: the assembly of a relevant protein structure dataset. This dataset assembly is the essential first step for using both distance-based methods included within these resources and character-based phylogenetic methods, which use 3Di structural characters to infer maximum likelihood trees [16] and helper tools like Structome-AlignViewer [17]. Thus, while these resources provide a direct route to distance-based trees, their fundamental purpose is to empower researchers with the curated data needed for any downstream structural evolutionary analysis.

Looking ahead, the data exploration process will be enhanced by incorporating advances in generative models. Future developments will explore the use of protein language model (pLM) embeddings as an alternative feature for structural comparison, alongside the planned integration of generative AI. This latter capability will enable automated textual summaries of search results, immediately highlighting common functional or taxonomic features within a structural neighbourhood. Such enhancements promise to transform this resource into an interactive knowledge discovery platform, accelerating insights into the evolution of protein structure and function.

## Availability and implementation

Structome-TM is available at https://biosig.lab.uq.edu.au/structometm/

